# Restraint of melanoma progression by cells in the local skin environment

**DOI:** 10.1101/2024.08.15.608067

**Authors:** Yilun Ma, Mohita Tagore, Miranda V. Hunter, Ting-Hsiang Huang, Emily Montal, Joshua M. Weiss, Yingxiao Shi, Tuulia Vallius, Peter K. Sorger, Richard M. White

## Abstract

Keratinocytes, the dominant cell type in the melanoma microenvironment during tumor initiation, exhibit diverse effects on melanoma progression. Using a zebrafish model of melanoma and human cell co-cultures, we observed that keratinocytes undergo an EMT-like transformation in the presence of melanoma, reminiscent of their behavior during wound healing. Surprisingly, overexpression of the EMT transcription factor Twist in keratinocytes led to improved overall survival in zebrafish melanoma models, despite no change in tumor initiation rates. This survival benefit was attributed to reduced melanoma invasion, as confirmed by human cell co-culture assays. Single-cell RNA-sequencing revealed a unique melanoma cell cluster in the Twist-overexpressing condition, exhibiting a more differentiated, less invasive phenotype. Further analysis nominated homotypic jam3b-jam3b and pgrn-sort1a interactions between Twist-overexpressing keratinocytes and melanoma cells as potential mediators of the invasive restraint. Our findings suggest that EMT in the tumor microenvironment may paradoxically limit melanoma invasion through altered cell-cell interactions.

## Introduction

The complex interplay between cancer cells and their microenvironment has emerged as a critical determinant of tumor progression and therapeutic response. In melanoma, the tumor microenvironment (TME) encompasses diverse cell types, including immune cells, fibroblasts, and endothelial cells^1^. However, during melanoma initiation the dominant cell type in the TME is the keratinocyte, an epithelial cell which makes up the majority of our skin surface. In normal homeostasis, each melanocyte reciprocally interacts with 30-40 keratinocytes^2^, and this interaction is essential for skin and hair color^3,4^. Despite decades of research, our understanding of keratinocytes in the context of melanoma remains incomplete.

Keratinocytes have been shown to both promote and inhibit tumor initiation. They are tightly adherent to melanocytes, and this can inhibit tumor development because nascent melanoma cells cannot escape the epidermis^5^. Previous work has demonstrated that newly initiated melanoma cells must escape from this adhesive network by loss of proteins such as PAR3^6^. In contrast, keratinocytes can also promote tumor development through secretion of growth factors such as endothelins or via GABAergic crosstalk between the two cell types^7,8^.

These conflicting data highlight that interactions between keratinocytes and nascent melanoma cells are likely dynamic and change rapidly during tumor initiation. Studying the nature of these interactions in human samples is challenging because biopsies are taken after the patient has come to the clinic, meaning that the earliest interactions in tumor initiation will be missed. This necessitates models which faithfully recapitulate the earliest stages of tumor initiation, yet have the cellular resolution to measure interactions between melanoma cells and keratinocytes.

In this study, we utilized a zebrafish model of melanoma to investigate the earliest interactions between melanoma cells and their neighboring keratinocytes^9^. Zebrafish have emerged as a powerful tool for cancer research due to their genetic tractability, conserved biology, and the ability to visualize tumor development and progression in real-time within the context of an intact organism^10,11^. Using a combination of cell-type specific genetic manipulations, in vivo imaging, and single-cell transcriptomics, we found that tumor-associated keratinocytes undergo changes associated with EMT, similar to what is found in wounded skin. Unexpectedly, we found that this keratinocyte EMT suppresses melanoma progression. This change in the keratinocytes occurs shortly after melanoma initiation, and results in keratinocytes which are more adhesive to these nascent tumor cells and prevents their movement out of the epidermis. Our data suggests that melanoma initiation revises an evolutionarily conserved wounding response in the nearby skin environment, which acts as a cell extrinsic tumor suppressor to prevent newly transformed cells from becoming fully tumorigenic.

## Results

### Melanoma initiation is associated with EMT in keratinocytes

To investigate the relationship between keratinocytes and melanoma cells *in vivo*, we created a transgenic zebrafish line in which GFP is expressed under the *krt4* promoter^12^. This line faithfully marks all adult keratinocytes present throughout the fish epidermis and scales, similar to previous lines using this promoter (Figure 1A). We then initiated melanomas in this background using the TEAZ method (Transgene Electroporation of Adult Zebrafish)^9,13^, in which plasmids containing oncogenes or sgRNAs against tumor suppressors can be introduced directly into the skin (Figure 1B). The major advantage of this method is that we can visualize melanoma initiation when the tumor is in its early stages, consisting of a small number of cells. We initiated tumors with a combination of BRAFV600E, sgRNAs against PTEN (*ptena/b)* and germline loss of p53 (Figure 1C). To account for skin wounding from electroporation in assessing changes in tumor-associated keratinocytes, we also performed TEAZ using a control vector that labels melanocyte-precursors but does not induce melanoma formation. Fluorescent imaging 8-weeks post-electroporation with the control vector demonstrated an injury-free epidermis, in contrast to the pronounced melanoma development in zebrafish administered with the oncogenic vectors (Figure 1D).

**Figure 1.**
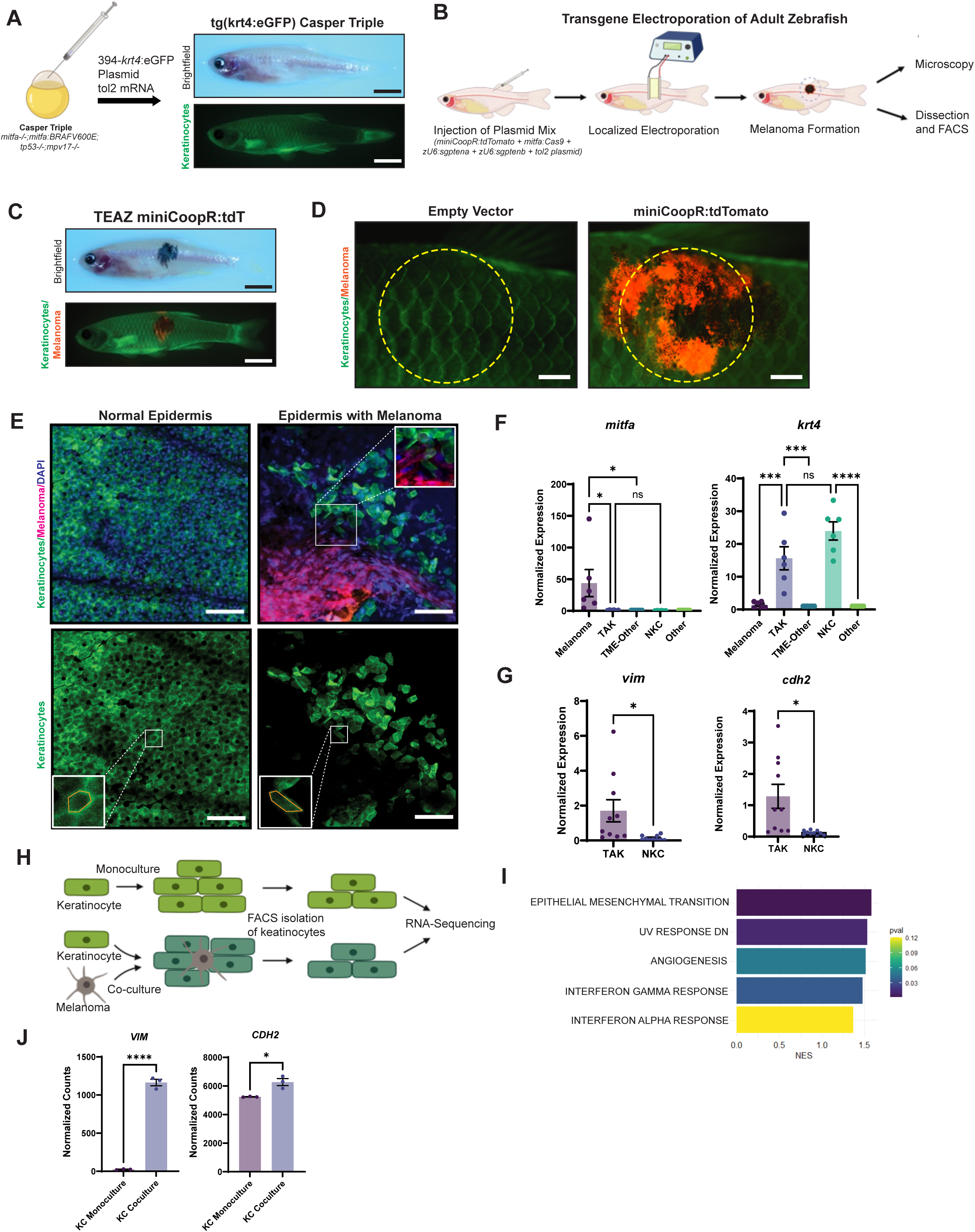
Keratinocytes in the melanoma microenvironment undergo EMT-like changes. (A) Generation of transparent zebrafish with GFP-labeling of keratinocytes. Casper Triples (*mitfa-/-;mitfa:*BRAFV600E;*tp53-/-;mpv17-/-*) were injected with the Tol2Kit 394 vector containing a *krt4*:eGFP cassette and tol2 mRNA. Brightfield shows a transparent zebrafish while fluorescence imaging shows eGFP-labeling of keratinocytes. (Scale bar = 5mm) (B) Schematic of TEAZ (Transgene Electroporation of Adult Zebrafish). Plasmid mix containing miniCoopR:tdTomato, *mitfa:*Cas9, *zU6:sgptena*, *zU6:sgptenb*, and the tol2 plasmid was injected superficially in the flank of the zebrafish. Electroporation of the injection site results in rescue of melanocyte precursors and the generation of a localized melanoma that could be analyzed by microscopy and FACS. (C) Brightfield and immunofluorescence of zebrafish 8-weeks post-TEAZ with localized and fluorescently labeled melanoma. (Scale bar = 5mm) (D) Immunofluorescence imaging of TEAZ region after 8-weeks, comparing empty vector control vs. miniCoopR:tdT conditions, with yellow dotted circles indicating general area of dissection for FACS. (Scale bar = 1mm) (E) Confocal imaging of zebrafish epidermis. Normal epidermis of Tg(*krt4:*eGFP) Casper Triple post-TEAZ with empty vector control shows eGFP-labeled, polygonal shaped keratinocytes regularly connected while epidermis with melanoma generated with *miniCoopR*-drived melanocyte rescue shows disrupted epidermis and irregularly shaped keratinocytes. Inset highlights melanoma cells with adjacent keratinocytes. (Scale bar = 50um) (F) qPCR of FACS sorted zebrafish epidermis with or without melanoma. tdTomato-labeled melanoma cells, eGFP-labeled keratinocytes and non-fluorescently labeled TME cells were isolated by dissection (as indicated in G) and FACS. Comparison of *mitfa* and *krt4* expression of samples normalized to non-fluorescent cells, either TME-Other in tumor samples or Other in non-tumor samples, shows enrichment of *mitfa* in melanoma sample and *krt4* in keratinocyte sample. ns = * = p<0.05, *** = p<0.001, **** = p<0.0001 by Tukey’s multiple comparisons test. (G) Comparison of the EMT-markers *vim (vimentin)* and *cdh2 (N-cadherin)* shows enrichment in TME keratinocytes vs. keratinocytes from epidermis without melanoma. * = p<0.05 by Welch’s t-test. (H) Schematic of keratinocyte-melanoma co-culture experiment. HaCaTs were cultured in monoculture or co-culture with A375 melanoma cells in triplicates for 21 days, followed by FACS isolation of keratinocytes for RNA-sequencing comparing co-culture vs. monoculture keratinocytes. (I) Top 5 enriched Hallmark pathways in HaCaTs co-cultured with A375 melanoma cells compared with HaCaTs in monoculture. (J) Normalized counts of EMT biomarkers vimentin (*VIM*) and N-cadherin (*CDH2*).

To better investigate the changes in the keratinocytes, we used confocal microscopy on fish 8 weeks after TEAZ. This revealed a marked disruption of keratinocyte morphology in the tumor bearing fish, which was not seen in control fish. We specifically noted disrupted cell-cell junctions, a disorganized pattern of keratinocytes, and loss of the normal hexagonal cell layer (Figure 1E). These changes were reminiscent of keratinocyte EMT, which has been previously noted to occur in wounded epidermis^14–16^. To further assess this possibility, we excised tissues from both tumor and control skin and used FACS to isolate keratinocytes (GFP+) and melanoma cells (tdTomato+) and performed qPCR. As expected, we found enrichment of *mitfa* in melanoma cells and *krt4* in keratinocytes, validating the successful cell-type isolation (Figure 1F). Comparative analyses between tumor-associated keratinocytes (TAKs) and normal keratinocytes (NKCs) from tissue without melanoma revealed upregulation of EMT markers vimentin and N-cadherin, consistent with our imaging results (Figure 1G).

We next asked whether these changes were also seen in human samples. To address this, we performed co-culture experiments between keratinocytes and melanoma cells. We grew GFP-labeled HaCaT keratinocytes either alone or with A375 melanoma cells for 21 days, followed by isolation by FACS for bulk RNA-sequencing (Figure 1H)^8^. Consistent with our *in vivo* results in the fish, the top pathway altered in the co-cultured HaCaT cells was enrichment of EMT (Figure 1I). Differential gene expression analysis showed notable upregulation of the mesenchymal markers vimentin and N-cadherin in co-cultured keratinocytes compared to monocultured control (Figure 1J), similar to what was found in vivo. Collectively, our data indicate that melanoma cells induce morphological and molecular markers of EMT in nearby keratinocytes.

### EMT transcription factors are upregulated in tumor-associated keratinocytes

EMT is usually driven by upstream transcription factors, which then act on downstream targets to repress adhesion molecules such as E-cadherin or activate other adhesion molecules such as N-cadherin^16^. We next wanted to understand which of these transcription factors was responsible for the EMT-like behavior in our keratinocytes. To address this, we utilized an existing scRNA-sequencing dataset of a BRAFV600E-driven zebrafish melanoma (Figure 2A). Dimensionality reduction with UMAP and subsequent clustering revealed two keratinocyte populations as indicated by module scoring for genes enriched in keratinocyte populations (Figure 2B-C). Subsequent differential gene expression and GSEA analysis of the two keratinocyte clusters revealed one cluster with enrichment for EMT, similar to what we observed above (Figure 2D-E). We refer to this EMT cluster as Tumor Associated Keratinocytes (TAKs), and the other cluster as a Normal Keratinocyte Cluster (NKCs). We focused on three EMT-transcription factors expressed in zebrafish keratinocytes, *snai1a*, *snai2*, *twist1a*, zebrafish homologs of human SNAIL, SLUG and TWIST. Differential expression showed significant enrichment in *snai1a* and *twist1a* in TAK versus NKC clusters (Figure 2F). The significant enrichment of *twist1a* in the TAK cluster, coupled with its rare expression in NKCs, positioned *twist1a* as a promising candidate for further investigation into its potential role in driving EMT-like changes in keratinocytes and, consequently, its impact on melanoma progression.

**Figure 2.**
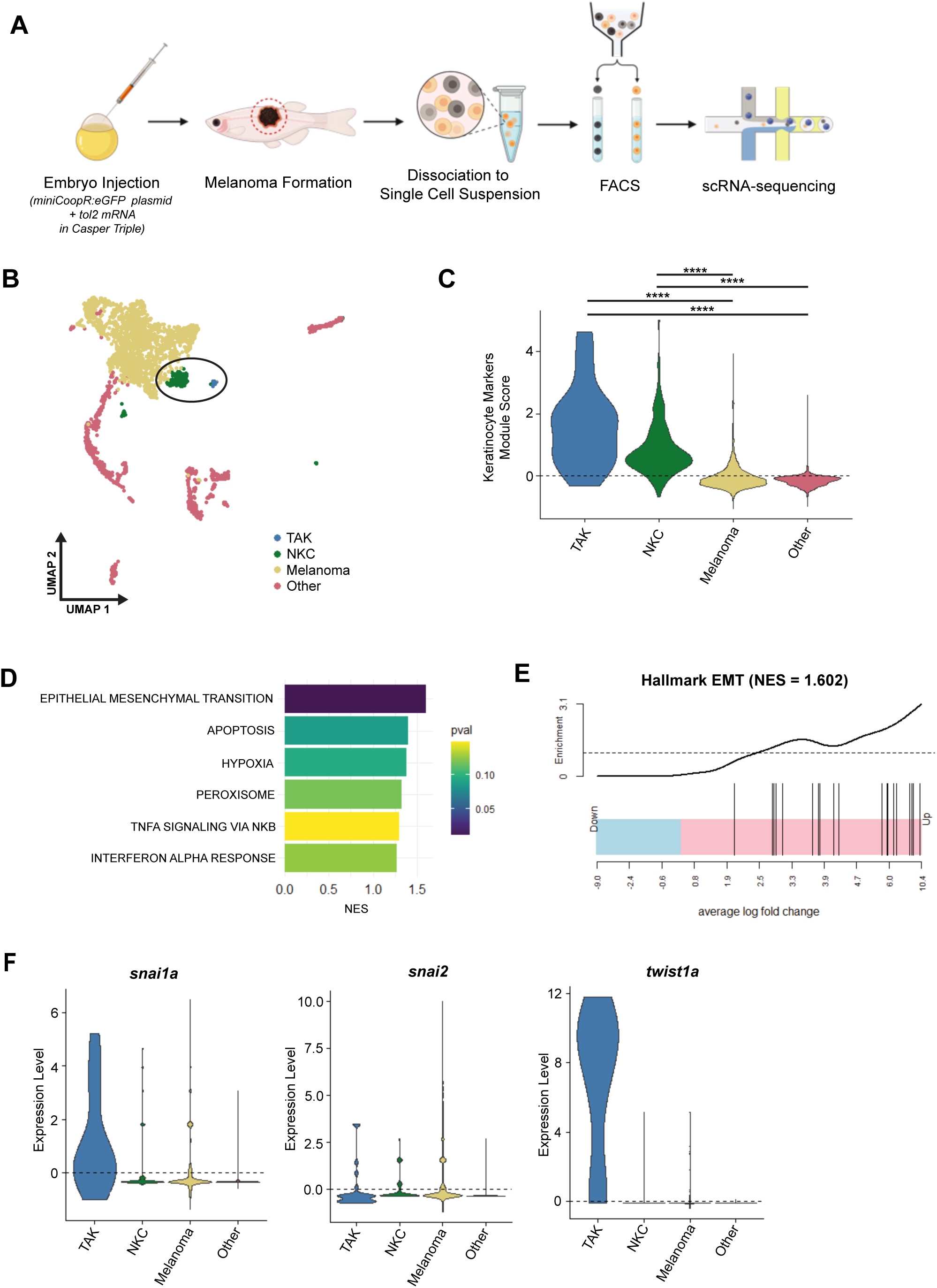
Zebrafish scRNA-sequencing shows upregulation of EMT-TFs in tumor-associated keratinocytes. (A) Schematic of scRNA-sequencing experiment. Embryo injection with miniCoopR:eGFP and tol2 mRNA in Casper Triple results in melanocyte rescue and subsequent melanoma formation. Melanoma was dissected and dissociated to single cell suspension for FACS isolation of eGFP+ melanoma cells and non-fluorescent TME cells for single cell RNA-sequencing. (B) UMAP highlighting two keratinocyte clusters, Melanoma and other TME cells. (C) Violin plot of keratinocyte module scores comparing between keratinocyte clusters, TAK and NKC, versus melanoma and other TME cells. (D) Top 6 GSEA Hallmark analysis comparing TAK vs. NKC (NES=Normalized Enrichment Score). (E) Hallmark EMT pathway enrichment in TAK vs NKC. (F) Enrichment of EMT-transcription factors in TAK vs. NKC, Melanoma, and Other TME cells.

### Keratinocyte *TWIST* restrains melanoma invasion in zebrafish

Having identified Twist as a potential driver of EMT-like changes in tumor-associated keratinocytes, we next asked how it affected melanoma phenotypes. We created new transgenic zebrafish in which the *krt4* promoter was used to drive the two zebrafish TWIST paralogs (*twist1a* and *twist1b*) in the context of BRAF-driven melanomas. In this experiment, we used standard 1-cell injection of the plasmids rather than TEAZ to ensure that the results were more generalizable to different melanoma initiation assays. Injected embryos were sorted at 5 days post-fertilization for tdTomato (melanocyte) and GFP (keratinocyte) expression, indicating successful melanocyte and keratinocyte transformation (Figure 3A). *These were mosaic animals, rather than stable lines, so we could directly assess the effect of TWIST on melanoma growth (although this precluded us from examining the effect of TWIST in keratinocytes without a nearby melanoma).* We monitored the fish for tumor-free survival as well as overall survival over the ensuing 26 weeks. Interestingly, we found no difference in melanoma initiation rate in the TWIST overexpression condition compared to empty vector (Figure 3B). Unexpectedly, however, we noted that overall survival was improved in the transgenic animals expressing TWIST in the keratinocytes (Figure 3B).

**Figure 3.**
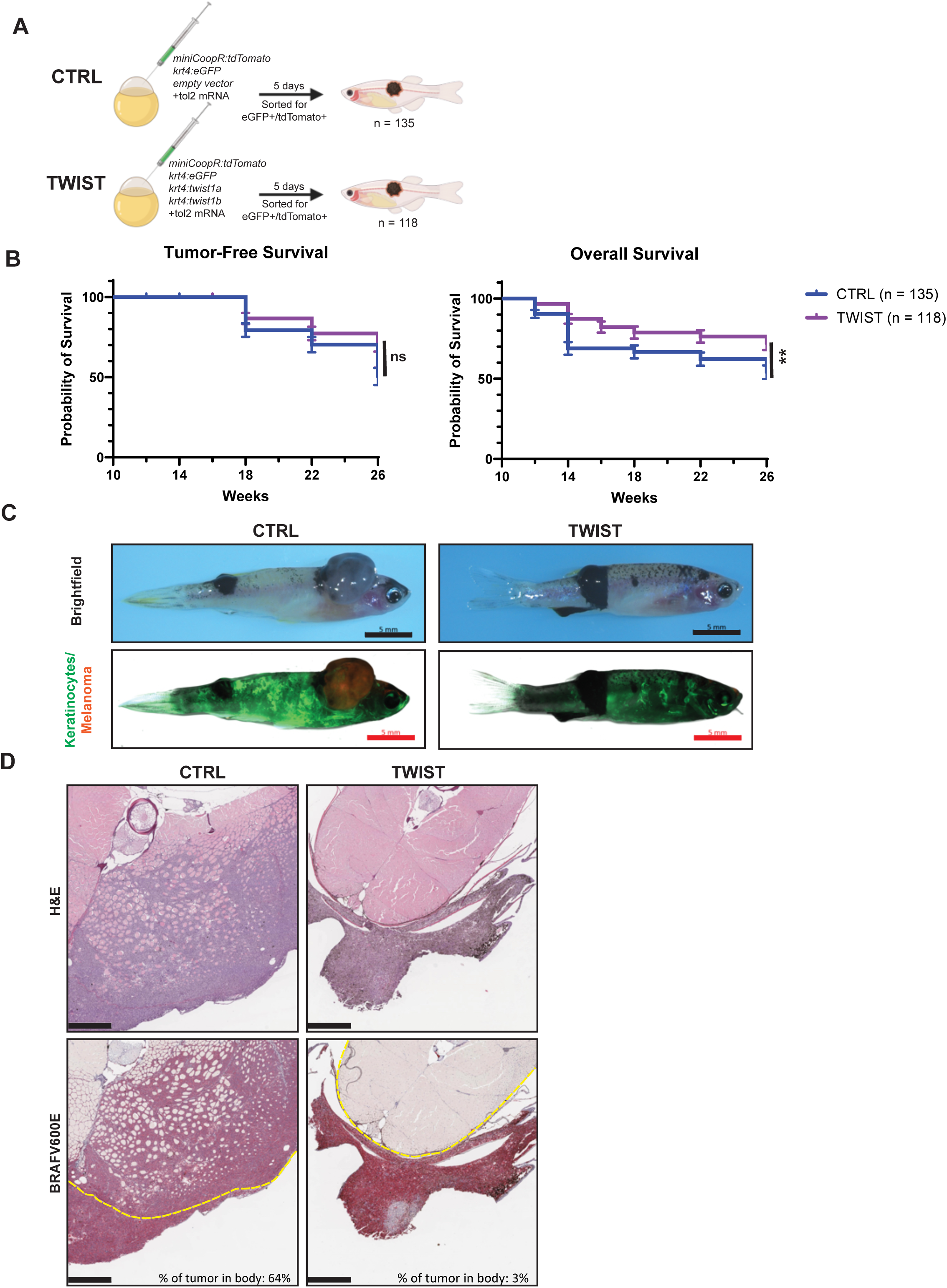
Overexpression of twist1a/b results in improved survival of fish with melanoma. (A) Schematic of zebrafish melanoma model with labeling and perturbation of keratinocytes. *Twist1a* and *twist1b* are overexpressed under the keratinocyte-specific, *krt4*, promoter in the TWIST condition and an empty vector control was used in the CTRL condition. Fish were sorted at 5 days for eGFP and tdTomato positivity as marker of successful keratinocyte labeling and melanocyte rescue (n = 135 CTRL, n = 118 TWIST). (B) Tumor-free survival and overall survival of A. ns = no significance, ** = p<0.005 by Log-rank (Mantel-Cox) test. (C) Sample images of zebrafish with melanoma at 26 weeks post-injection. Keratinocytes are labeled by GFP, melanoma are labeled by tdTomato. Melanomas are pigmented. Scale bar = 5mm. (D) H&E and IHC of cross-sections through zebrafish body and melanoma. Dotted yellow line demarcates border of body. % of tumor in body is calculated as tumor area within body border divided by total tumor area. Scale bar = 500um.

This discrepancy between melanoma-free and overall survival suggested that the tumors in the TWIST condition should be phenotypically distinct from the control tumors. To assess this, we performed immunohistochemistry on the tumors *(n=3 in each condition)* and surrounding tissues from the control and TWIST conditions (Figure 3C). The oncogenic driver in this melanoma model is hBRAFV600E and serves as an IHC marker for the tumor cells. Comparison of hBRAFV600E staining revealed significant melanoma infiltration into the zebrafish body in the CTRL condition, as opposed to a nearly non-invasive tumor in the TWIST condition (Figure 3D). The lack of melanoma invasion was also observed when the melanoma developed in other anatomical locations as well (Supp. Figure 1), although because survival started to decrease around 26 weeks, we could not let these fish grow longer to see if they eventually became invasive at very late time points. These findings suggest that *twist1a/twist1b* overexpression in keratinocytes does not impair tumor initiation, but instead impairs melanoma invasion and improves survival when expressed in the microenvironment.

Because this was a somewhat unexpected finding, we wanted to ensure that this was not an artifact of the zebrafish system. We therefore developed an assay to measure interactions between human keratinocytes and melanoma cells. To simulate melanoma invasion into neighboring cells, we developed a novel cell culture-based invasion assay using HaCaT lines and multiple melanoma cell lines. Melanoma cells cultured on poly-L-lysine coated coverslips were placed atop keratinocytes, allowing assessment of melanoma cell infiltration after 20 hours (Figure 4A). Due to the migratory nature of melanoma cells, they will migrate off the coverslip and infiltrate the layer of keratinocytes, allowing us to assess relative differences between culture conditions.

**Figure 4.**
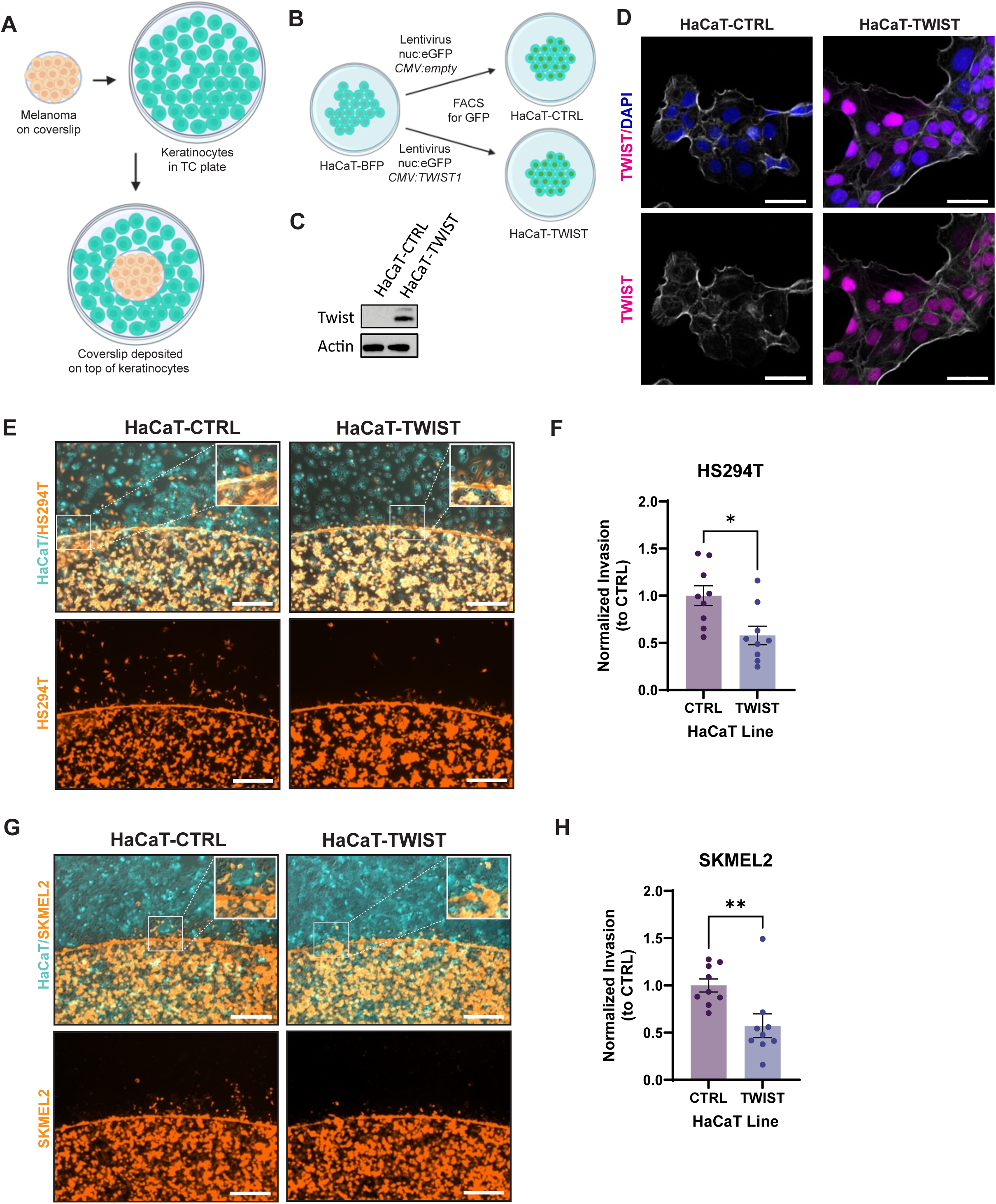
Zebrafish findings are recapitulated in human cell lines. (A) Schematic of coverslip cell infiltration assay. Melanoma cells are plated on a coverslip and allowed to attach overnight. The coverslip is then transferred into a well of keratinocytes to assess melanoma infiltration into keratinocytes. (B) Generation of a HaCaT cell line overexpressing TWIST1. HaCaT-BFP was infected with lentivirus containing cassette with nuclear localized GFP and CMV driven TWIST1 or no ORF. Infected cell lines were allowed to grow for a week before sorting for nuclear GFP as a marker of successful integration. (C) Western blot for Twist expression in HaCaT-CTRL and HaCaT-TWIST. (D) Immunofluorescence imaging for Twist localization in HaCaT-CTRL and HaCaT-TWIST. TWIST staining is pseudo-colored in magenta, DAPI in blue, phalloidin in white. Scale bar = 50um. (E) Immunofluorescence imaging of coverslip cell infiltration assay with HS294T-tdT (orange) melanoma cells in co-culture with either HaCaT-CTRL or HaCaT-TWIST (cyan). Insets highlight areas of melanoma cell infiltration into keratinocyte monolayers. Scale bar = 500 μm. (F) Quantification of E. Infiltrating HS294T melanoma cells from each image were counted and averaged across four images per well. Resulting cell counts were normalized to average cell counts of HaCaT-CTRL from each set. N = 9, 3 sets, 3 replicates/wells per set. * = p<0.05 by t-test. (G) Immunofluorescence imaging of coverslip cell infiltration assay with SKMEL2-tdT (orange) melanoma cells in co-culture with either HaCaT-CTRL or HaCaT-TWIST (cyan). Insets highlight areas of melanoma cell infiltration into keratinocyte monolayers. Scale bar = 500 μm. (H) Quantification of G. Infiltrating SKMEL2 melanoma cells from each image were counted and averaged across four images per well. Resulting cell counts were normalized to average cell counts of HaCaT-CTRL from each set. N = 9, 3 sets, 3 replicates/wells per set. * = p<0.05 by t-test.

HaCaT keratinocytes were transformed via lentiviral infection to overexpress TWIST1 (HaCaT-TWIST) or an empty vector control (HaCaT-CTRL) (Figure 4B). Western blot analysis confirmed robust Twist protein overexpression in HaCaT-TWIST compared to HaCaT-CTRL (Figure 4C), and immunofluorescence imaging revealed nuclear localization of Twist (Figure 4D). Co-culture of HS294T melanoma cells with HaCaT-TWIST resulted in significantly reduced melanoma cell invasion into keratinocytes compared to HaCaT-CTRL (Figure 4E-F), similar to what we had observed above in our zebrafish system. This finding was recapitulated using SKMEL2, demonstrating the inhibitory effect of TWIST1 overexpression in keratinocytes on melanoma invasion across different cell lines (Figure 4G-H). However, it was also possible that it was something lacking in the TWIST keratinocytes that led to less melanoma invasion (rather than actively suppressing invasion). To test this, we also performed a control assay in which we allowed tdTomato melanoma cells to invade into unlabeled melanomas of the same line. Using both HS294T and SKMEL2 cells, these cells do have some migratory capacity in this setting (Supp. Figure 2). This assay cannot discern whether the TWIST keratinocytes actively suppress invasion versus lack of something that promotes invasion, which will be an area for future studies. Collectively, our results demonstrate that induction of EMT in keratinocytes is associated with reduced melanoma invasion and improvement in animal survival.

### Keratinocyte EMT promotes aberrant adhesion to nascent melanoma cells

While EMT within the tumor cell is well recognized to promote invasion, our data suggest that EMT in the microenvironment paradoxically restrains tumor invasion. We wanted to better understand the downstream mechanisms accounting for this result. We therefore analyzed our control vs. TWIST tumors using single-cell RNA sequencing, which would allow us to understand potential mechanisms by which melanoma cells were interacting with these keratinocytes. Melanoma and adjacent skin from three fish per condition (CTRL or TWIST) were dissociated into single cell suspensions and FACS-sorted for eGFP+ keratinocytes and tdTomato+ melanoma cells. To enrich the keratinocyte population, keratinocyte and melanoma cell suspensions were combined at a 7:3 ratio for each condition and prepared for scRNA-sequencing (Figure 5A). The resulting CTRL and TWIST datasets were filtered for quality control as described (Supp Figure 3A). UMAP dimensional reduction of the integrated dataset shows distinct clusters of melanoma and keratinocytes, with high eGFP expression in keratinocyte clusters and high tdTomato expression in melanoma clusters (Figure 5B). Furthermore, we identified two KC clusters in the present in both the CTRL and TWIST conditions and a third cluster of KCs in the TWIST condition. We performed differential gene expression (DEG) analysis between clusters and GSEA on differentially expressed genes between the two clusters present in both conditions showed enrichment in EMT as seen in figure 2, identifying EMT-enriched cluster as TAK and other as NKC (Supp Figure 3B). As expected, we also identified a unique cluster of KCs only present in the TWIST dataset that highly expresses *twist1a/b* (Supp Figure 3C), the genes that we overexpressed in the TWIST condition, and labeled this cluster Twist-High (Figure 5C).

**Figure 5.**
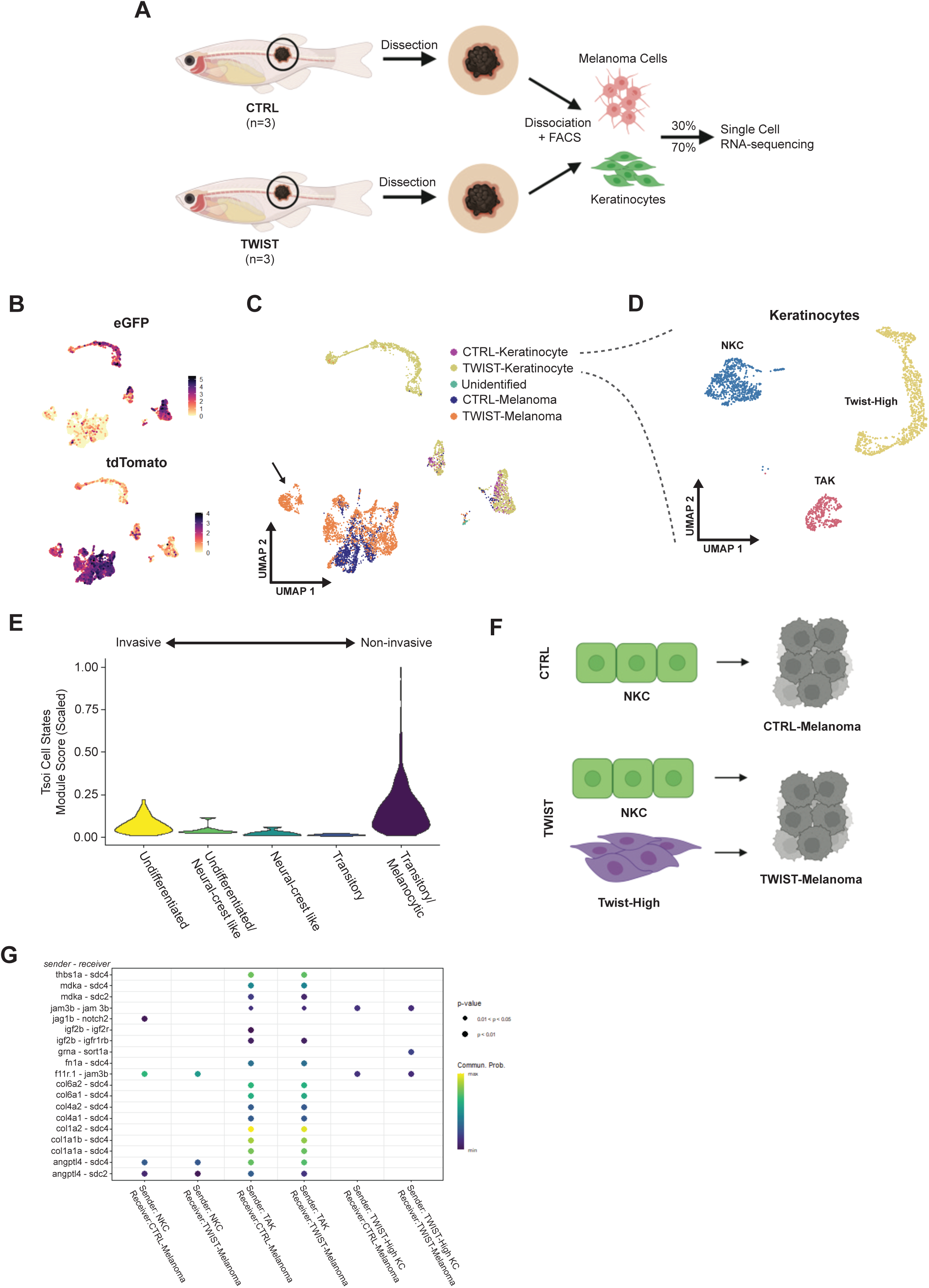
scRNA-sequencing shows unique keratinocyte-melanoma communication with twist1a/b overexpression in keratinocytes. (A) Schematic of scRNA-sequencing protocol. Melanoma and surrounding tissue were dissected from 26-weeks old zebrafish from either CTRL or TWIST conditions as shown in Figure 3. Samples were dissociated to single cell suspensions for FACS isolation of keratinocytes (GFP) and melanoma (tdTomato). Keratinocytes and melanoma were recombined per condition at a ratio of 7:3 for enrichment of keratinocytes for scRNA-sequencing. (B) UMAP dimensional reduction and feature plots of scRNA-sequencing dataset. CTRL and TWIST samples were sequenced, and cell types were identified using eGFP+ for keratinocytes and tdTomato+ for melanoma, after which the datasets were integrated. (C) UMAP of keratinocyte clusters from CTRL and TWIST conditions combined, showing Normal Keratinocyte Cluster (NKC), Tumor-Associated Keratinocyte (TAK), and Twist-High cluster (TWIST-specific). CTRL: 32.9% TAK. TWIST: 14.3% TAK, 56.3% Twist-High. (D) UMAP highlighting keratinocyte clusters, with a Tumor-Associated Keratinocyte (TAK) cluster, a Normal Keratinocyte Cluster (NKC), and a Twist-High cluster unique to the TWIST condition. (E) Melanoma cell state analysis of the melanoma cluster unique to the Twist condition indicated by arrow in Figure 5C. (F) Schematic overview of CellChat analysis. In CTRL condition, we analyzed Ligand-Receptor pairs with NKC as sender and CTRL-Melanoma as receiver. In TWIST condition, we analyzed L-R pairs with both NKC and Twist-High as sender and TWIST-Melanoma as receiver. (G) CellChat analysis results. L-R pairs shown at p<0.01, with color scale indicating communication probability of L-R pair.

We first analyzed the melanomas that arose in the control vs. TWIST animals by comparing their gene signature to well-annotated signatures of human melanoma cell states. It is now widely recognized that melanomas exist in at least 4 different transcriptional states, ranging from undifferentiated/invasive, to neural crest, to intermediate, to melanocytic/proliferative^17^. These distinct states are mutually exclusive, yet exhibit a high degree of plasticity, with interconversion between the states^17^. *Interestingly, we found a cluster of melanoma cells that developed only in the TWIST condition was enriched for the melanocytic/proliferative state but not undifferentiated/invasive gene markers (Figure 5E). Consistent with this, Gene Set Enrichment Analysis (GSEA) of the TWIST condition also showed significant enrichment for oxidative phosphorylation (Supplemental Figure 3D-F), which we previously showed is representative of the melanocytic state*^18^*. Because this state is amongst the least invasive ones, this is consistent with our in vivo observations that these melanomas are phenotypically less invasive*.

We hypothesized that this change in cell state might be induced by physical interactions between the TWIST keratinocytes and the melanoma cells. To address this, we analyzed potential cell-cell interactions using CellChat, a software tool that allows us to quantitatively characterize and visualize cell-cell communications using a curated zebrafish ligand-receptor interaction database^19^ (Figure 5F). Two unique ligand-receptor pairings were identified that only occur between TWIST keratinocytes and the melanomas that arose in these animals: a homotypic *jam3b-jam3b* interaction and a *pgrn-sort1a* (progranulin-sortilin) interaction (Figure 5G).

The *jam3b* interaction was of particular interest to us, as this protein has been recently identified as one required for melanophore survival in zebrafish^20^ and for human melanoma metastasis^21,22^. In normal human skin, JAM3 is expressed in keratinocytes of the superficial epidermis, whereas its heterotypic partner JAM1 is only expressed in basal keratinocytes^23,24^. Since melanomas largely arise in this basal area, this suggests that TWIST expression in the keratinocytes was leading to aberrant expression of jam3b, forming stronger homotypic jam3b-jam3b attachment between these keratinocytes and the melanoma cells and preventing their invasion.

### Conservation in human studies

Finally, we wished to determine if the TWIST^hi^ state in our zebrafish models was also present in human melanomas. We use the GeoMX spatial transcriptomics platform to interrogate a series of early melanoma precursor lesions, a subset of a larger study using this method to look at spatial organization of melanoma stages^25^. We first stained sections with antibodies to cytokeratin (to mark keratinocytes) and SOX10/MART1 (to mark melanocytes and melanoma cells). This allows for selection of microregions of interest (MR) which are then subject to RNA-seq (Figure 6A-D). From this, we then queried for the TAK/TWIST gene sets found in the zebrafish. Whereas normal skin melanocytes showed no such enrichment, there was a enrichment of these signatures as the lesions progressed from early to late precursor (Figure 6E). This suggests that EMT-like alterations in keratinocytes are an early event in both zebrafish and human melanoma.

**Figure 6.**
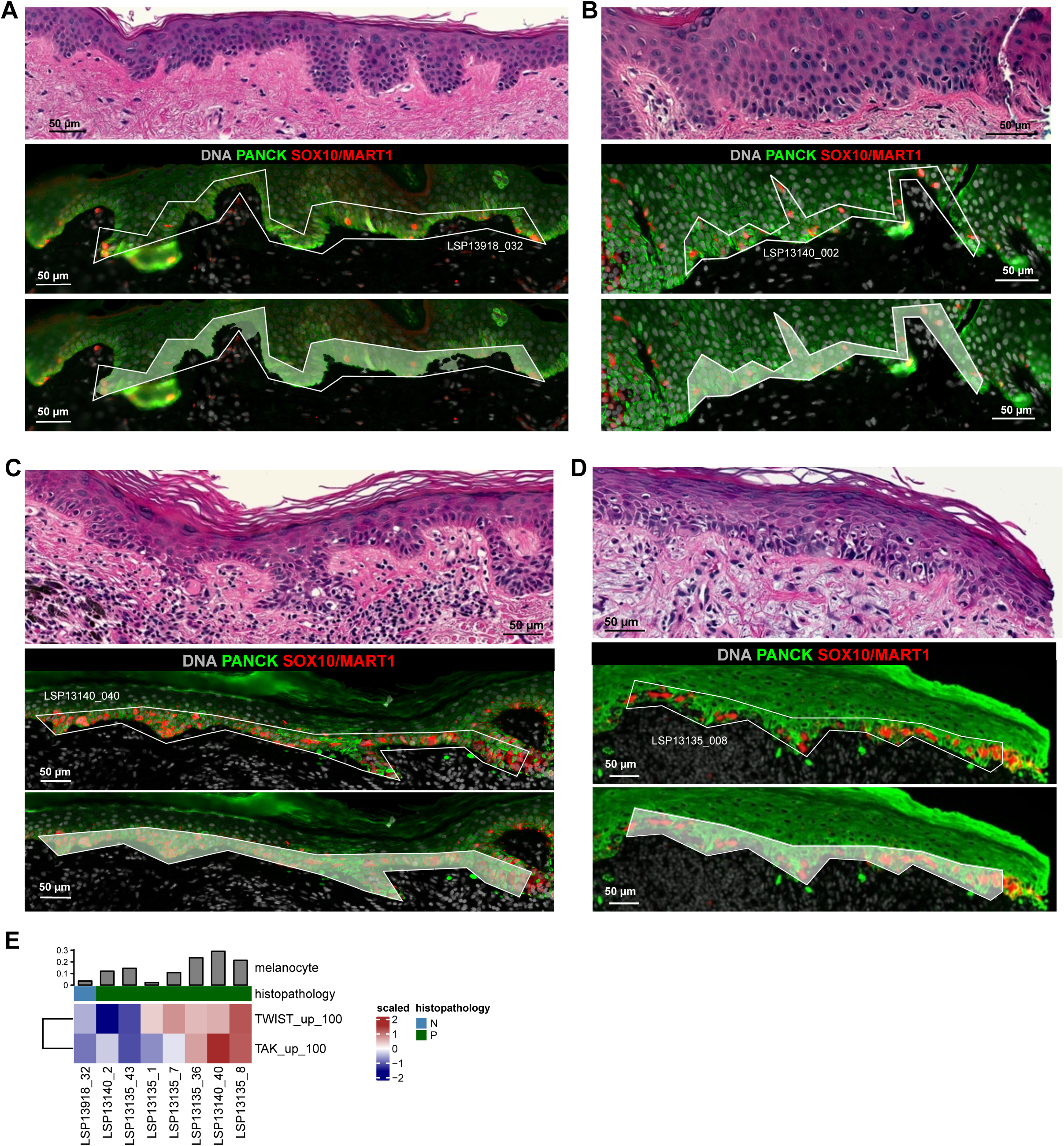
Enrichment of keratinocyte signatures in microregions with melanocytic atypia. (A-D) Visualization of a subset of reanalyzed GeoMx microregions (MRs); GeoMx data (n=8 MRs) is reanalyzed from Vallius, Shi, Novikov, et al. biorxv 2025. Immunofluorescence staining is shown for morphology markers DNA (grey), panCK (green) and SOX10/MART1 (red; middle and bottom rows). The selected MRs are illustrated with white polygons (middle), and the shaded area within the MRs represent the area for oligonucleotide barcode collection (referred as a “segmentation mask” per NanoString; bottom). Serial sections were H&E-stained and the regions mapping to the sequenced MRs are shown (top row). The MRs were annotated as epidermis containing morphologically normal melanocytes (panel A), and regions of melanocytic atypia (precursor; panels B-D). Scale bars, 50 μm. (E–G) Melanocyte-adjusted TAK and TWIST gene-signature activity across reanalyzed MRs (n=8 GeoMx MRs). (E) Heatmap showing gene-signature scores derived from the top 100 upregulated genes defining the TAK and TWIST programs across MRs. Each column represents an MR, and the MRs are annotated for sample ID and histopathological stage, with melanocyte fraction (melanocyte count divided by total nuclei count within the MR) per GeoMx MR displayed as a top annotation.

## Discussion

In this study *(summarized in Figure 7)*, we observed that keratinocytes in a zebrafish of melanoma undergo an EMT-like transformation in the presence of melanoma. This alteration is reminiscent of keratinocyte behavior during wound healing, in which keratinocytes exhibit markers and morphological changes associated with EMT in development^15,26^. Interestingly, we also observed an increase in N-cadherin expression in KC, which is usually attributed to melanoma as it becomes more aggressive and invades into the dermis to associate with fibroblasts. Our findings would suggest that if keratinocytes are also upregulating N-cadherin expression, their normal contact-based regulatory controls on melanoma may still be relevant, though not acting through E-cadherin. Additionally, re-analysis of a published zebrafish melanoma scRNA-sequencing dataset showed distinct populations of keratinocytes that expressed markers of EMT, demonstrating the feasibility of studying this KC population using our zebrafish model and nominating Twist1 as a potent EMT-TF in this cell ^27^. Twist expression has been found to be upregulated at the edge of wounded skin upon treatment with bFGF, a well-characterized growth factor in melanoma^26,28^. If melanoma acts as an open-wound in the skin, then it is possible that Twist1 might be upregulated in tumor-associated keratinocytes (TAKs) to close this wound.

**Figure 7.**
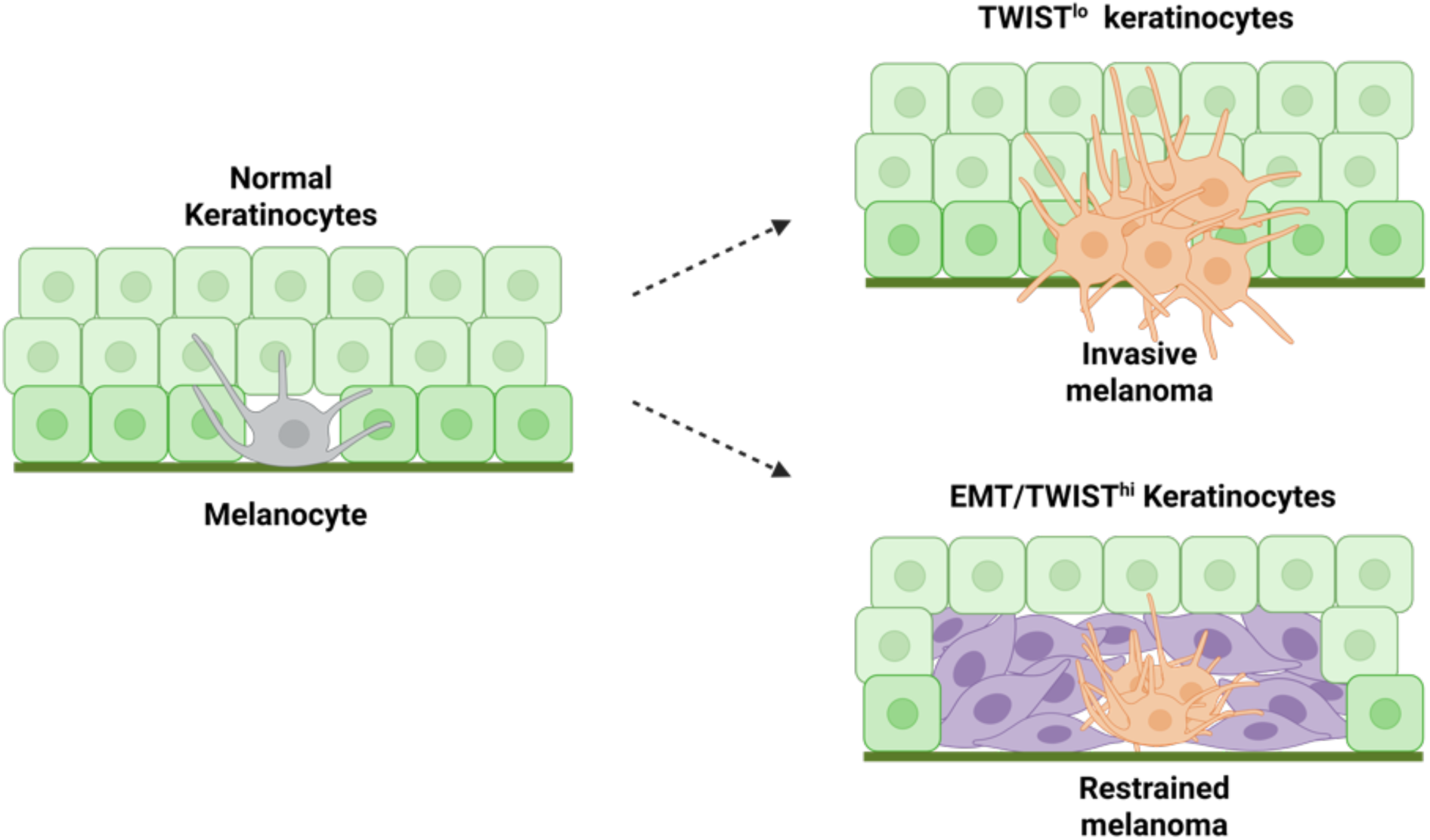
Model of melanoma restraint by keratinocytes. Schematic model of melanoma-keratinocyte interactions in the skin. During early melanoma development, melanocytes reside within an epidermis composed of normal keratinocytes. In the context of low TWIST expression (TWIST^lo^ keratinocytes), melanoma cells adopt an invasive, undifferentiated phenotype and are able to breach the epidermal compartment. In the presence of EMT/TWIST^hi^ keratinocytes, melanoma cells are restrained within the epidermis and adopt a more differentiated, less invasive cell state. These findings suggest that EMT-like changes in tumor-associated keratinocytes, rather than promoting tumor spread, act as a cell-extrinsic barrier to melanoma invasion through altered cell-cell interactions in the local skin microenvironment.

Our zebrafish melanoma survival experiment showed improvements in overall survival in zebrafish with KCs overexpressing Twist compared to those that received an empty vector, despite the fish forming tumors at the same rate. This survival improvement was shown to be caused by a decrease in melanoma invasion, raising the possibility that Twist overexpressing KCs could restrain melanoma invasion. Human cell co-cultures with a HaCaT cell line overexpressing Twist showed a similar finding to our in vivo zebrafish model, with reduced melanoma cell infiltration into keratinocytes. However, an important caveat to these cell culture experiments is that the keratinocyte densities vary when co-cultured with different melanoma lines (i.e. Hs.294t vs. SKMel2), the reasons for which are not known. However, because we saw a consistent pattern across these two lines, it is less likely that our results are due to these differences in keratinocyte density alone.

To learn more about the dynamics of melanoma and TME KCs, we performed scRNA-sequencing on our zebrafish melanoma model to account for both KCs in contact with melanoma and those in the periphery. This yielded three KC clusters in fish overexpressing Twist, while two clusters in fish that received the empty vector; accounting for KCs overexpressing Twist and the KC populations we had observed previously in our reanalysis of a zebrafish melanoma scRNA-sequencing dataset^27^.

Interestingly, we found a cluster of melanoma cells unique to the TWIST condition, which shared gene signatures similar to that of the genes observed in both transitory and melanocytic states as published by Tsoi et al.^17^. These cell states were defined to be more differentiated with higher MITF expression and correlated to a more proliferative but less invasive cohort from Hoek et al.^29^. Thus, it is likely that this melanoma cluster unique to the TWIST condition could be responsible for the reduced overall melanoma invasion.

Further analysis using CellChat nominated pgrn-sort1a and jam3b-jam3b as unique interactions between Twist-High KCs and TWIST-Melanoma cells. As previously described, jam3b has been identified as a critical protein in zebrafish melanophore survival with known involvement in melanoma metastasis. The aberrant expression of jam3b on Twist-overexpressing KCs could indicate strong homotypic interaction with melanoma jam3b that retains the melanoma in the epidermis. Although the progranulin-sortilin interaction has not been characterized in melanoma, sortilin has been identified as a key regulator of progranulin levels^30^. Progranulin is known to be constitutively expressed by KCs, which could be cleaved to epithelins that promote KC proliferation^31,32^. Progranulin is also a potent mediator of the wound response produced by dermal fibroblasts in addition to epidermal keratinocytes^33^. Perhaps the increase in progranulin is cleared by melanoma cells through sortilin, resulting in the endocytosis and lysosomal transport of sortilin^34^. The degradation of sortilin could be responsible for decreased cell migration and invasion, as sortilin is required for the interaction of proNGF, a neurotrophin produced by melanoma, with NGFR in promoting melanoma migration^35^. Further studies are needed to elucidate the precise mechanisms underlying the nominated interactions, jam3b-jam3b and pgrn-sort1a, and to explore their potential as therapeutic targets in melanoma.

## Methods

### Zebrafish husbandry

Zebrafish were maintained in a dedicated facility with controlled temperature (28.5 °C) and salinity. The fish were kept on a 14-hour light/10-hour dark cycle and fed a standard zebrafish diet consisting of brine shrimp followed by Zeigler pellets. Embryos were obtained through natural mating and incubated in E3 buffer (5 mM NaCl, 0.17 mM KCl, 0.33 mM CaCl2, 0.33 mM MgSO4) at 28.5 °C. For procedures requiring immobilization, zebrafish were anesthetized using Tricaine-S (MS-222, Syndel) prepared as a 4 g/L stock solution with a pH of 7.0. The stock solution was protected from light exposure and diluted to the appropriate concentration to achieve fish immobilization. All experimental procedures and animal protocols described in this manuscript were conducted in compliance with the Institutional Animal Care and Use Committee (IACUC) protocol #12-05-008, approved by the Memorial Sloan Kettering Cancer Center (MSKCC).

### Generation of zebrafish line with fluorophore labeled keratinocytes

Embryos at the one-cell stage from the Casper Triple zebrafish line (mitfa:BRAFV600E;p53-/-;mitfa-/-;mpv17-/-)^9,36^ were injected with a krt4:eGFP expression cassette in the 394 vector of the Tol2Kit^37^ with tol2 mRNA. Larvae were sorted for positive GFP fluorescence at day three and raised to adult for breeding. F0 fish were in-crossed and resulting F1 were outcrossed with Casper Triple zebrafish for consistent GFP expression. Starting from F2, the krt4:eGFP zebrafish line was maintained by out-crossing with Casper Triple zebrafish and sorting for GFP expression.

### Transgene Electroporation in Adult Zebrafish (TEAZ)

TEAZ was utilized to generate melanoma as previously described^9,38^. Krt4:eGFP zebrafish (krt4:eGFP Casper Triple) were anesthetized with tricaine and injected with a plasmid solution containing miniCoopR-tdT (250ng/μl), mitfa:Cas9 (250 ng/μl); zU6:sgptena (23 ng/μl) with target sequence GAATAAGCGGAGGTACCAGG, zU6:sgptenb (23ng/μl) with target sequence GAGACAGTGCCTATGTTCAG, and the tol2 plasmid (55ng/μl). For control electroporation without generating melanoma, zebrafish were injected with mitfa:tdTomato (250ng/μl), mitfa:Cas9 (250 ng/μl), zU6:non-targeting (46ng/μl) and tol2 plasmid (55ng/μl). All fish were injected on the left flank below the dorsal fin and electroporated with the BTX ECM 830 electroporator using 3mm platinum Tweezertrodes (BTX Harvard Apparatus; #45-0487). Electroporator settings used: LV Mode, 40V, 5 pulses, 60ms pulse length, and 1s pulse interval. Electroporated zebrafish were screened for successful electroporation 7 days post-electroporation by tdTomato expression using fluorescence microscopy and melanoma tracked by imaging once per week. All live zebrafish imaging were performed with the Zeiss AxioZoom V16 fluorescence microscope.

sgRNA sequences for TEAZ listed below:

Nontargeting: AACCTACGGGCTACGATACG

ptena: GAATAAGCGGAGGTACCAGG

ptenb: GAGACAGTGCCTATGTTCAG

### Confocal imaging of zebrafish epidermis

Zebrafish with or without melanoma were anesthetized in tricaine (MS-222, Syndel) as described above. Site of injection is visually identified by the presence of melanoma or the area below the dorsal fin. Scales were removed with tweezers and fixed in 4% PFA in PBS (Santa Cruz 281692) in a 96-well plate for 15 minutes. Fixed scales were washed three times with PBS and permeabilized with 0.1% Triton-X 100 (Thermo Scientific 85111) in PBS, then blocked with 10% goat serum (Thermo Fisher 50062Z). Scales were incubated with 1:250 GFP polyclonal antibody, Alexa Fluor 488 (Thermo Fisher A21311) overnight at 4°C. Next day, scales were washed three times with PBS, incubated with 1:1000 Hoechst 33342 (Thermo Fisher H3570) for 1 hour and mounted onto slides with VECTASHIELD Vibrance Antifade Mounting Media (Vector Laboratories H-1700). Samples were imaged on the Zeiss LSM880 inverted confocal microscope and images were processed using FIJI v1.53.

### Flow cytometry of adult zebrafish cells

Zebrafish were euthanized using ice-cold water. Melanoma and adjacent skin were dissected from fish with melanoma and skin alone was dissected from below the dorsal fin. Subsequently, samples were cut into 1mm strips using a clean scalpel and placed into 15 mL conical tubes (Falcon 352099) with 3 ml of DPBS (Gibco 14190250) and 187.5μl of 2.5 mg/ml Liberase TL (Roche 5401020001). Samples were incubated in dissociation solution at room temperature for 30 minutes on a shaker with gentle movement to prevent tissue from settling at the bottom of the tube. At 15 minutes of incubation, a wide bore p1000 pipette tip (Thermo Scientific 2079G) was used to gently pipette the sample up and down for 90 seconds. After 30 minutes, 250 μl of FBS (Gemini Bio) was added to stop the enzymatic activity of Liberase TL and samples were pipetted up and down using a wide bore p1000 pipette tip for 90 seconds. Dissociated cells were then filtered through a 70 μm cell strainer (Falcon 352350) into a 50 mL conical tube (Falcon 352098) placed on ice. Samples were centrifuged at 500g at 4°C for 5 minutes and supernatant was removed by pipetting. The cell pellet was resuspended in 500 μl of PBS with 5% FBS and filtered again through 40 μm tip filters (Bel-Art H136800040) into 5 ml polypropylene tubes (Falcon 352063). For subsequent FACS analysis, 0.5 μl of 1000x DAPI (Sigma-Aldrich D9542) was added to each sample. Samples were FACS sorted (BD FACSAria) at 4°C for GFP-positive keratinocytes and tdTomato-positive melanoma gated using fluorophore-negative zebrafish controls.

### Zebrafish tissue RNA extraction and real-time quantitative PCR (RT-qPCR)

FACS sorted zebrafish cells were deposited directly into 750 μl TRIzol LS Reagent (Invitrogen 10296010) in Eppendorf DNA LoBind Tubes (Eppendorf 022431021). After collection, samples were snap-frozen using dry ice and stored at −80°C. RNA extraction was performed per TRIzol LS manufacturer protocols. For precipitation of RNA, 10ug supplemental glycogen (Roche 10901393001) was used per sample to account for low cell numbers. Resulting RNA was resuspended in Nuclease-free water (Fisher Scientific AM9937). 25ng RNA per sample was transcribed to cDNA using Superscript III First-Strand Synthesis System (Invitrogen 18080051). cDNA mix was diluted 1:10 with Nuclease free water for RT-qPCR using Power SYBR Green PCR Master Mix (Applied Biosystems 4368708) and the Bio-Rad CFX384 Touch Real-Time PCR System (Bio-Rad 1855484). Resulting Cq values were normalized to *hatn10* as previously described.

qPCR primer sequences:

*hatn10* fwd: 5’-TGAAGACAGCAGAAGTCAATG-3′

*hatn10* rev: 5′-CAGTAAACATGTCAGGCTAAATAA-3′

mitfa fwd: 5’-GGCACCATCAGCTACAATGA-3’

mitfa rev: 5’-GAGACAGGGTGTTGTCCATAAG-3’

krt4 fwd: 5’-GGAGGTGTTTCCTCTGGTTATG-3’

krt4 rev: 5’-GAACCGAATCCTGATCCACTAC-3’

vim fwd: 5’-GGATATTGAGATCGCCACCTAC-3’

vim rev: 5’-GACTCTCGCAGGCTTAATGAT-3’

cdh2 fwd: 5’-GAGCCATCATCGCCATACTT-3’

cdh2 rev: 5’-CTTGGCCTGTCTCTCTTTATCC-3’

### scRNA-sequencing Analysis of Miranda et al

Zebrafish scRNA-seq data from ref^27^ was re-analyzed using R 4.2.0. and Seurat 4.3.0^39,40^. Cluster identities were maintained as published. Keratinocyte Module Scores were calculated using the AddModuleScore function with default parameters using published gene lists. Differential Gene Expression (DGE) analyses between clusters were performed using FindMarkers. Differentially expressed gene lists were converted from zebrafish genes to human orthologs using DIOPT as previously described^41,42^. GSEA analysis on differentially expressed genes between keratinocyte clusters was performed using fgsea 1.22.0 and the Hallmark pathways set from MSigDB^43,44^.

### Twist overexpression in zebrafish keratinocytes

*Twist1a* (ENSDART00000043595.5) and *twist1b* (ENSDART00000052927.7) were TOPO cloned into the attL1-L2 Gateway pME vector and LR cloned into the pDestTol2pA2 vector (Tol2Kit 394) with p5E-*krt4* promoter and p3E-polyA (Tol2Kit 302). To generate the zebrafish melanoma model as previously described, one-cell stage Casper Triple zebrafish embryos (mitfa:BRAFV600E;p53-/-;mitfa-/-;mpv17-/-) were injected with miniCoopR-tdTomato, krt4-eGFP, either krt4-twist1a and krt4-twist1b for Twist overexpression condition or empty vector for control condition, tol2 mRNA and phenyl red. Injections were performed three times on different days with parents from the same clutch. Embryos were grown at standard conditions and sorted at 5 days post-injection for eGFP and tdTomato expression using the Zeiss AxioZoom V16 fluorescence microscope. eGFP+/tdTomato+ fish in the CTRL (n=135) and TWIST (n=118) conditions were maintained to adulthood.

### Zebrafish imaging and tumor-free survival tracking

Zebrafish were regularly monitored for melanoma formation and survival every 4 weeks, beginning at 10 weeks post-fertilization. Melanoma formation was screened visually using the Zeiss AxioZoom V16 fluorescence microscope under 20X magnification. Kaplan-Meier curves and corresponding statistics were generated using GraphPad Prism 9. Statistical differences in survival between conditions were determined by the Mantel-Cox log-rank test.

### Histology of zebrafish samples

Zebrafish were euthanized in tricaine (MS222, Syndel). Each fish was dissected in three sections consisting of head, body, and tail. Samples were placed in 4% PFA in PBS (Santa Cruz 281692) for 72h on a shaker at 4°C, then paraffin embedded. Histology was performed by HistoWiz Inc. (histowiz.com) using a Standard Operating Procedure and fully automated workflow. Samples were processed, embedded in paraffin, and sectioned at 5μm. Immunohistochemistry was performed on a Bond Rx autostainer (Leica Biosystems) with enzyme treatment (1:1000) using standard protocols. Sections were stained with H&E or IHC with antibodies including BRAFV600E (ab228461) and GFP (ab183734). Bond Polymer Refine Detection (Leica Biosystems) was used according to the manufacturer’s protocol. After staining, sections were dehydrated and film coverslipped using a TissueTek-Prisma and Coverslipper (Sakura). Whole slide scanning (40x) was performed on an Aperio AT2 (Leica Biosystems).

### Cell culture

Human melanoma lines A375, HS294T, and SKMEL2 were obtained from ATCC. Human keratinocyte line HaCaT was obtained from AddexBio. All cells were routinely tested and confirmed to be free from mycoplasma. Cells were maintained in a humidified incubator at 37°C and 5% CO2. Cells were maintained in DMEM (Gibco 11965) supplemented with 10% FBS (Gemini Bio) and split when confluent, approximately 2-3 times per week.

### Twist overexpression in HaCaT

The HaCaT cell line was labeled with eBFP to allow for identification during co-culture. 293T (ATCC) was transfected with the pLV-Azurite plasmid (Addgene 36086) with pMD2.5 (Addgene 12259) and psPAX2 (Addgene 12260) using Invitrogen Lipofectamine 3000 Transfection Reagent (Invitrogen L3000015) according to manufacturer protocol. HaCaTs were infected with lentivirus containing CMV:eBFP and selected for eBFP positivity using ampicillin and FACS. Subsequently, HaCaT-eBFP was infected with lentivirus containing CMV:TWIST1 (Horizon Precision LentiORF Human TWIST1 OHS5898-202622685) or CMV:empty control created by removing ORF of CMV:TWIST1 plasmid. HaCaT line overexpressing TWIST1 was labeled as HaCaT-TWIST and HaCaT line with empty vector was labeled as HaCaT-CTRL. HaCaT lines were subsequently sorted for nuclear eGFP expression present in the plasmid as part of the Precision LentiORF system. HaCaT-CTRL and HaCaT-TWIST were cultured as previously described.

### Western blot

Cells were washed with DPBS (Gibco 14190250) and lysed in RIPA buffer (Thermo Scientific 89901) with the addition of protease and phosphatase inhibitors (Thermo Scientific 78440). Lysates were centrifuged at 13,000g at 4 °C and quantified using the Pierce BCA Protein Assay Kit (Pierce 23227). Samples were reduced with the laemmli SDS-sample buffer (Boston BioProducts BP111R) and boiled for 10 minutes. For gel electrophoresis, samples were loaded into 4-15% precast protein gels (Bio-Rad 4561084), then transferred to 0.2µm nitrocellulose membranes (Bio-Rad 1704158). Membranes were washed in TBST and blocked with 5% milk in TBST (Boston BioProducts P1400) for 1 hour at RT. Membranes were washed and incubated with primary antibodies overnight. Antibodies used includes Twist (Abcam ab50887) and beta-actin (CST 3700S). On the next day, membranes were washed with TBST and incubated with appropriate secondary antibodies for 1 hour. Blots were incubated with the Immobilon Western Chemiluminescent HRP Substrate (Millipore WBKLS0500) and imaged with the Amersham ImageQuant 800.

### Immunofluorescence

Cells were cultured on chamber slides (Thermo Scientific 154739) overnight at standard cell culture conditions. Culture media was washed with DPBS (Gibco 14190250) and fixed with 2% PFA in PBS (Santa Cruz 281692) for 15 minutes at RT. Cells were washed with DPBS and permeabilized with 0.1% Triton-X 100 (Thermo Scientific 85111) in PBS, then blocked with 10% goat serum (Thermo Fisher 50062Z) for 1 hour at RT. Primary antibodies used include Twist (Abcam ab50887). Cells were incubated with primary antibody in 10% goat serum overnight at 4 °C and washed with DPBS the next day, before incubation with 1:1000 Hoechst 33342 (Thermo Fisher H3570) for 1 hour and mounted with VECTASHIELD PLUS Antifade Mounting Medium (Vector Laboratories H1900). Slides were imaged on the Zeiss LSM880 inverted confocal microscope and images were processed using FIJI v1.53.

### Melanoma infiltration assay

RFP-labeled melanoma cell lines, including A375, SKMEL2, HS294T, were plated on poly-l-lysine coated round glass coverslips (Corning 354085) placed in 24-well plates at 150-200k cells per coverslip. HaCaT cell lines were plated in 6-well plates at 250-300k cells per well. Cells were allowed to attach overnight and the coverslip containing melanoma cells is transferred to 6-wells containing HaCaT cell lines using tweezers. All coverslips were placed in the center of the well. KC-melanoma co-cultures were incubated in standard cell culture conditions for 24 hours. Co-cultures were imaged by fluorescence microscopy at 4 locations of each coverslip: top, right, bottom and left, to capture variations in melanoma cell infiltration into KC lines. FIJI v1.53 was used to count the number of infiltrating melanoma cells per image and average infiltrating melanoma cells were calculated per well. All experiments were performed in 3 sets, with 3 replicates per set per condition. Average infiltrating cell numbers per well were normalized to the average infiltrating cell number per well in the HaCaT-CTRL condition.

### scRNA-sequencing analysis of zebrafish melanoma

Six zebrafish, three each from CTRL and TWIST conditions at 26 weeks post-injection were selected for scRNA-sequencing of melanoma tumors. To account for set differences, one fish from each of three injection sets were chosen in each condition. Melanoma and adjacent skin were dissected from the fish, then enzymatically and mechanically dissociated into single cell solutions as described above. The samples were FACS sorted for GFP and RFP positivity, corresponding to eGFP expressed by keratinocytes and tdTomato expressed by melanoma. The sorted cells were placed in (Gibco 11965) supplemented with 10% FBS (Gemini Bio) and 1% penicillin-streptomycin-glutamine. To enrich for KCs, sorted KCs and melanoma cells from each fish were recombined at a 7:3 KC:melanoma ratio. Sorted cells were pelleted and resuspended in DPBS + 0.1% BSA. Samples were also combined based on their genetic perturbation condition. Droplet-based scRNA-seq was performed using the Chromium Single Cell 3′ Library and Gel Bead Kit v3 (10X Genomics) and Chromium Single Cell 3′ Chip G (10X Genomics). 10,000 cells were targeted for encapsulation. GEM generation and library preparation was performed according to kit instructions. Libraries were sequenced on a NovaSeq S4 flow cell. Resulting reads were aligned to the GRCz11 reference genome with the addition of eGFP and tdTomato sequences using CellRanger v5.0.1 (10x Genomics). scRNA-sequencing analysis was performed as detailed above. In addition, melanoma cells were scored using AddModuleScore to assess their enrichment of genes associated with the four main melanoma cell states and intermediate states^17^. The highest scoring gene module for each cell was annotated as its cell state. CellChat^19^ was used to analyze cell-cell communication between KC and melanoma clusters using its zebrafish L-R database.

### Spatial transcriptomics (GeoMx) data reanalysis

The GeoMx spatial transcriptomics data was reanalyzed from Vallius et al, biorxiv 2025^25^. First, the reanalyzed MRs were selected based on the availability of the keratinocyte compartment within the sequenced MRs. Only the MRs with keratinocytes present within the profiled region (referred as the “segmented area” per NanoString; examples shown in Fig 6 panels A-D) were reanalyzed and included in this manuscript. The H&E-stained serial sections were used for histopathological annotation of the samples, as described in Vallius et al. The biospecimen metadata and GeoMx MR annotations are shown in Supplementary Table 1. To account for the variable cell type composition across MRs, we first computed the melanocyte fraction for each MR as the ratio of melanocyte count to total nuclei count. The melanocyte count was determined by the SOX10/MART1 immunofluorescence staining, by visually counting the nuclei staining positively for SOX10 and/or MART1. A non-melanocyte scaling factor was then defined as 1−melanocyte fraction. With the top 100 upregulated genes in the TAK and TWIST programs, the raw gene-signature scores were calculated using Gene Set Variation Analysis (GSVA, RRID:SCR_021058) and subsequently normalized by dividing each score by the corresponding non-melanocyte scaling factor, yielding keratinocyte-adjusted signature values. For visualization, keratinocyte-adjusted signature scores were assembled into a matrix (signatures × MRs) and standardized by row (z-score scaling across MRs) to emphasize the relative differences between MRs. Heatmaps were generated using the ComplexHeatmap package in R (RRID:SCR_017270), with columns representing samples and rows representing gene signatures. Rows were hierarchically clustered, while columns were ordered using optimal leaf ordering (OLO) based on pairwise sample distances.

### Statistics and reproducibility

Statistical analysis and figures were generated by GraphPad Prism 9, R Studio 4.2.0 and Biorender.com. Image processing was performed in FIJI v1.53. Statistical tests are described in figure legends and methods. Experiments were repeated at least three times unless otherwise noted. All animal and cell experiments were performed with a reasonable number of replicates by power calculations or feasibility of the experimental method. GeoMx gene expression data^25^ is available via the Gene Expression Omnibus (GEO; RRID:SCR_005012).

---

**Supplemental Figure 1.**
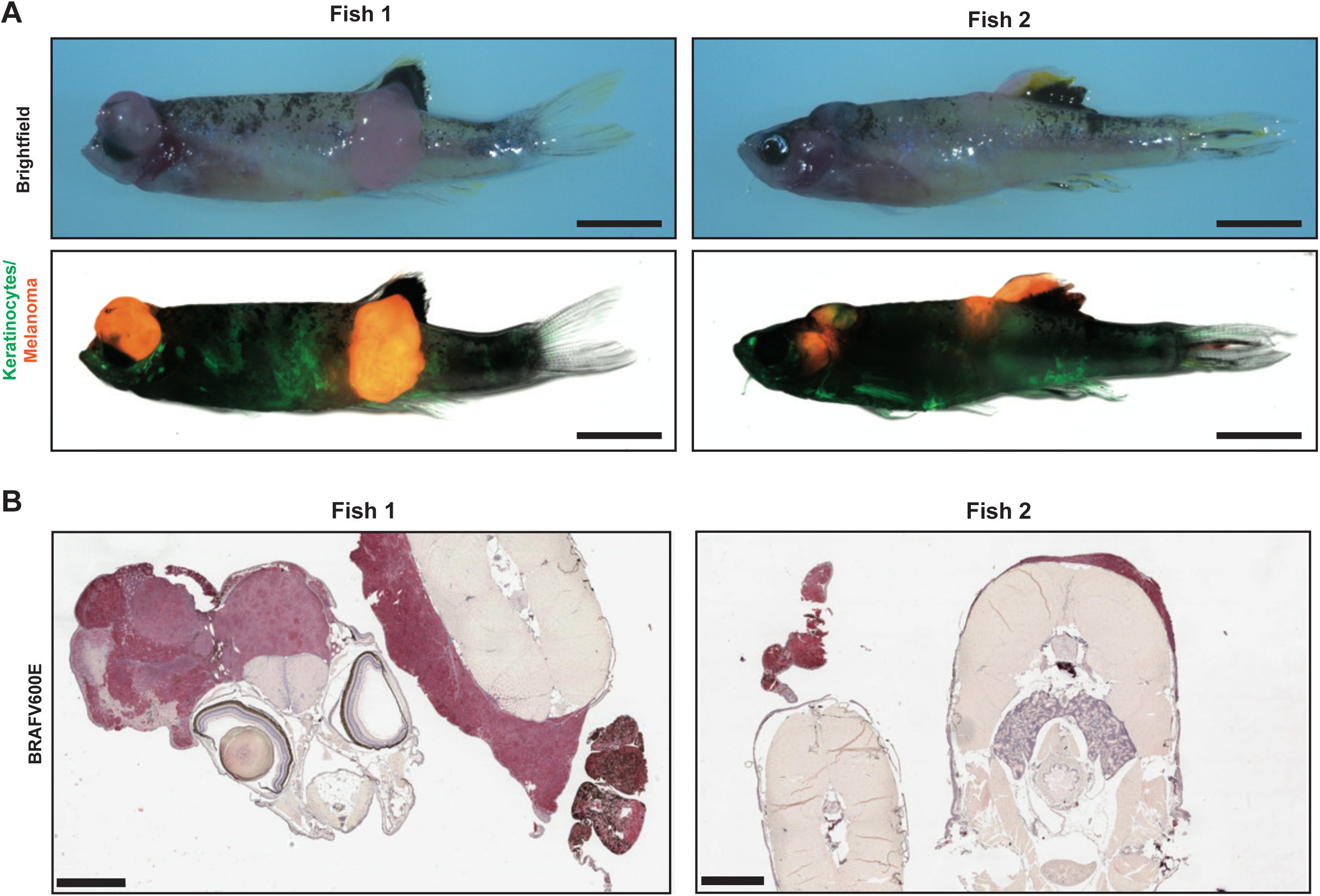
Melanoma in TWIST conditions were non-invasive independent of anatomical location. (A) Brightfield and fluorescence images of two sample fish from the TWIST condition. Melanoma are pigmented in brightfield images. Keratinocytes are labeled by eGFP and melanoma are labeled by tdTomato in fluorescence images. Scale bar = 5mm. (B) IHC of cross-sections through the zebrafish body and melanoma. Red staining indicates positivity for hBRAFV600E. Scale bar = 1mm.

**Supplemental Figure 2.**
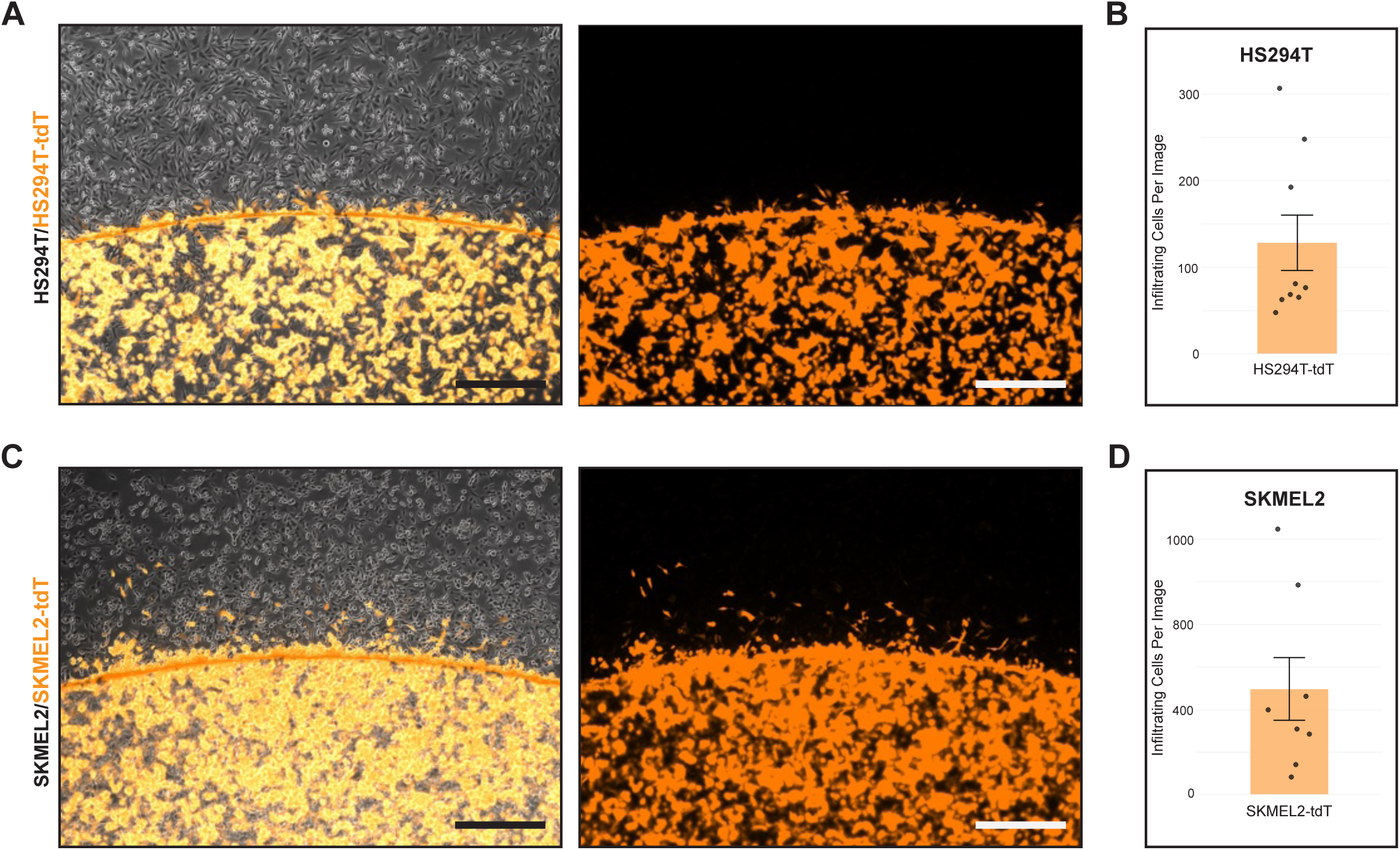
Melanoma migration from coverslip. (A) Immunofluorescence imaging of coverslip cell infiltration assay with HS294T-tdT (orange) melanoma cells in co-culture with parental HS294T (uncolored). Scale bar = 500um. (B) Quantification of the migration of HS294T-tdTomato melanoma cells in co-culture with parental HS294T melanoma cells. Each data point is the sum of 4 images per well. Mean ± SE, n = 9 wells from 3 independent experiments. (C) Immunofluorescence imaging of coverslip cell infiltration assay with SKMEL2-tdT (orange) melanoma cells in co-culture with parental SKMEL2(uncolored). Scale bar = 500um. (D) Quantification of the migration of SKMEL2-tdTomato melanoma cells in co-culture with parental SKMEL2 melanoma cells. Each data point is the sum of 4 images per well. Mean ± SE, n = 8 wells from 3 independent experiments.

**Supplemental Figure 3.**
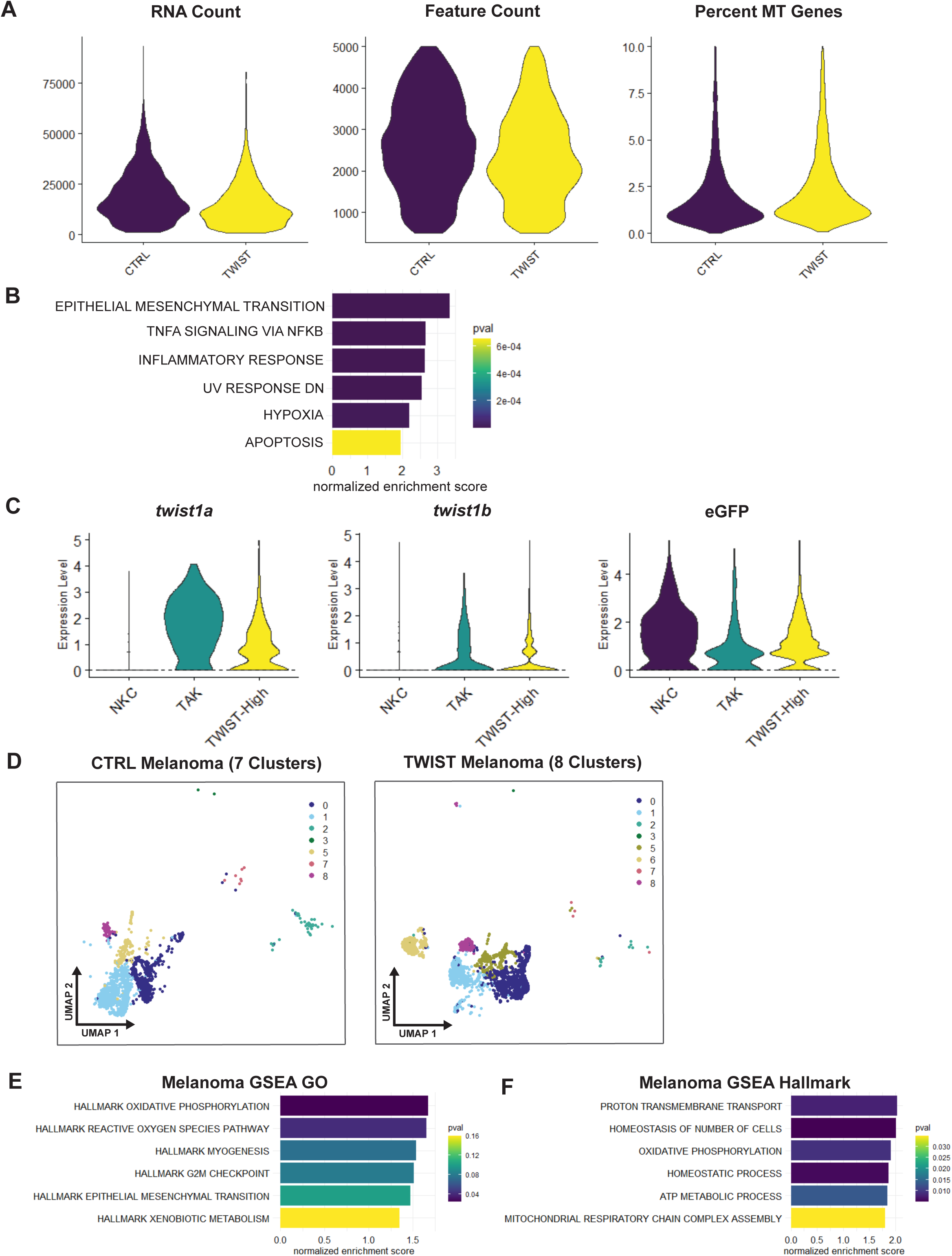
scRNA-sequencing of zebrafish melanoma. (A) Violin plots of scRNA-sequencing data comparing CTRL and TWIST conditions after quality control. Samples were filtered by number of features between 500 and 5000, with less than 10% mitochondria genes per cell. (B) Top 6 GSEA Hallmark pathways of differentially expressed genes between TAK and NKC clusters. (C) *Twist1a*, *twist1b*, and eGFP expression in keratinocyte clusters. (D) UMAP of melanoma samples from either the CTRL or TWIST conditions, identified by seurat clusters. 7 distinct clusters were identified in the CTRL melanoma sample vs. 8 distinct clusters in the TWIST melanoma sample. (E-F) GSEA analysis showing pathways upregulated in TWIST vs. CTRL melanoma: (E) GO:Biological Process pathways and (F) Hallmark pathways.

